# Obesity and Tumor Development Reprogram the Proteome and Metabolic Effects of Adipose- and Tumor-Derived Extracellular Vesicles

**DOI:** 10.64898/2026.02.08.704727

**Authors:** Emma J. Grindstaff, Laith A. Rayyan, Yazan Alwarawrah, Erika Rezeli, Nancie J. MacIver, Dorothy Teegarden, Stephen D. Hursting, Ximena Bustamante-Marin

## Abstract

Obesity alters systemic metabolism and immune function, yet how obesity and tumor progression regulate extracellular vesicle (EV) composition and function within the tumor microenvironment remains unclear. Using a preclinical model of diet-induced obesity (DIO) and triple-negative breast cancer (TNBC), we investigated how obesity and tumor stage shape the proteomic composition of EVs from visceral adipose tissue (VAT-EVs) and mammary tumors (tumor-EVs), and how these EVs regulate immune and tumor cell metabolism. Orthotopically transplanted metM-Wnt^lung^ tumors were classified as early (∼0.5 cm³) or late (∼1.0 cm³), and EV proteomes were analyzed by mass spectrometry.

At early stages, tumor-EVs from DIO mice, compared with control lean mice, were depleted in immune-related proteins, whereas VAT-EVs were enriched in mitochondrial and fatty acid oxidation proteins. In contrast, at later stages, tumor-EVs from DIO mice were enriched in lipid metabolism and oxidative stress–associated proteins, while VAT-EVs exhibited loss of mitochondrial proteins consistent with metabolic dysfunction. Functionally, tumor-EVs and VAT-EVs differentially regulated CD8 T cell mitochondrial activity and cytokine production and induced distinct, stage-dependent metabolic reprogramming in non-aggressive epithelial-like (E-Wnt) versus mesenchymal-like (M-Wnt) tumor cells.

These findings suggest that obesity and tumor progression dynamically reshapes EV cargo, enabling EV-mediated metabolic reprogramming that may contribute to immune suppression and TNBC progression.

## Introduction

Obesity is a chronic metabolic disease characterized not only by excess body weight but also by visceral (abdominal) adiposity, a metabolically active fat depot that promotes insulin resistance, adipokine imbalance, chronic inflammation, and vascular dysfunction, thereby creating a tumor-permissive microenvironment (Chooi et al., 2019, Trevellin et al., 2023) (Engin, 2017b). Obesity is a well-established risk factor for breast cancer incidence, recurrence, and mortality, particularly after menopause (Engin, 2017a, Harbeck et al., 2019, Bustamante-Marin et al., 2021) (Reeves et al., 2007). Growing evidence indicates that central adiposity—often measured by waist circumference or waist-to-hip ratio—is specifically associated with estrogen receptor–negative breast cancer, including triple-negative breast cancer (TNBC) (Dietze et al., 2018, Pierobon and Frankenfeld, 2013). TNBC disproportionately affects younger, frequently premenopausal women and lacks expression of estrogen receptor, progesterone receptor, and HER2, limiting targeted therapeutic options (Obidiro et al., 2023, Millikan et al., 2008). Consequently, women with visceral obesity who develop TNBC experience poorer clinical outcomes, including higher risks of recurrence, metastasis, and mortality (Dent et al., 2007).

The complex communication between visceral adipose tissue (VAT), breast epithelial cells, and immune cells plays a critical role in mediating the detrimental effects of obesity on breast cancer progression (Alhallak et al., 2021, Fletcher et al., 2018, Hillers-Ziemer et al., 2021). In this context, EVs have emerged as key mediators of intercellular communication (Lago-Baameiro et al., 2025, Ziglari et al., 2025, Wang et al., 2025). EVs are membrane-bound vesicles released from the plasma membrane or endosomal compartments, carrying a diverse cargo of nucleic acids, proteins, and lipids (Li et al., 2021). They can interact with recipient cells by binding to surface receptors or by delivering their contents directly into the cytoplasm (Yáñez-Mó et al., 2015). Due to their heterogeneity in size, composition, and biogenesis, EVs have garnered significant interest as regulators of cellular function (Sun and Chang, 2024, Maas et al., 2017)

Tumor cells exploit EVs to communicate with both neighboring and distant cells, modulate the tumor microenvironment and facilitate processes such as extracellular matrix remodeling and metabolic reprogramming to support invasion and metastasis (Njock et al., 2022, Xu et al., 2024, Ma et al., 2025, Lee et al., 2025). Given their central role in cell signaling, EVs are being explored for their potential as vehicles for therapeutic delivery and as sources of diagnostic and prognostic biomarkers of cancer, including TNBC (Das et al., 2023, Haney et al., 2020). However, their role in the interplay between obesity and breast cancer across different stages of tumor progression remains unclear. Here, using a preclinical model of diet-induced obesity (DIO) and triple-negative breast cancer (TNBC), and employing tumor volume as a proxy for tumor stage (early vs. late), we tested the hypothesis that obesity and tumor progression reprogram the proteomic composition of tumor- and VAT–derived EVs, thereby altering immune- and metabolism-related proteins to promote immune suppression and tumor progression.

### Experimental Procedures

#### Mammary Cancer Cell Lines

The mammary TNBC cell lines used in this study (E-Wnt, M-Wnt, and metM-Wnt^lung^) were developed in the Hursting laboratory from MMTV-Wnt1 transgenic mouse model of basal-like/triple-negative breast cancer on a C57Bl/6 background. E-Wnt and M-Wnt represent epithelial-like and mesenchymal-like, non-metastatic tumor phenotypes within the same genetic background, whereas metM-Wnt^lung^ is a metastatic derivative selected for robust tumor formation and EV production in vivo (O’Flanagan et al., 2017, Dunlap et al., 2012). All cell lines were cultured in RPMI1640 media without phenol red (GIBCO Life Technologies) supplemented with 10% FBS, 10 mmol/L HEPES buffer, 2 mmol/L L-glutamine, and 1% penicillin/streptomycin.

#### Mouse Model of Obesity and TNBC

All animal procedures were approved by the University of North Carolina Institutional Animal Care and Use Committee. We used a previously established mouse model of diet-induced obesity (DIO) and TNBC (Zheng et al., 2013, O’Flanagan et al., 2017), and the overall experimental design is summarized in Figure 1A. Briefly, 30 C57BL/6 female mice (7 weeks of age) were allowed to acclimate for 1 week, being fed a control diet (10 kcal% fat, D12450J, Research Diets, New Brunswick, NJ) ad libitum. Then they were randomly assigned to one of two dietary groups, each fed ad libitum for study duration (n=15/group): control diet to induce a lean phenotype (CON), or high-fat diet (60 kcal% fat, sucrose-matched to control diet, D12492, Research Diets) to induce an obese phenotype (DIO). After 20 weeks on diet (when obesity was well established in the DIO group), all mice were orthotopically engrafted in the fourth mammary gland fat with 50,000 metM-Wnt^lung^ cells. Within each dietary group, the mice were further randomized into 2 subgroups based on target tumor volume, which in this study was used as an experimental proxy for tumor stage: 1) mice bearing early tumors, in which the tumor was collected when it reached approximately 0.5 cm^3^ and classified as “early” (CON-0.5 and DIO-0.5; n = 7, each) and 2) mice bearing late tumors were those with a tumor volume of approximately 1 cm^3^ and classified as “late” (CON-1 and DIO-1, n = 8. each). This classification was based solely on tumor size, a key parameter in the TNM system, but does not represent full clinical staging, which also considers nodal involvement and metastasis (Amin et al., 2017). Tumor growth was monitored and measured daily. The mice were euthanized humanely with CO_2_ when the tumor reached the target volume. Mammary tumor and VAT (including periovarian and mesenteric fat) were collected from the same mouse (samples were not pooled), weighed and stored at -80 °C for future processing.

**Figure 1:**
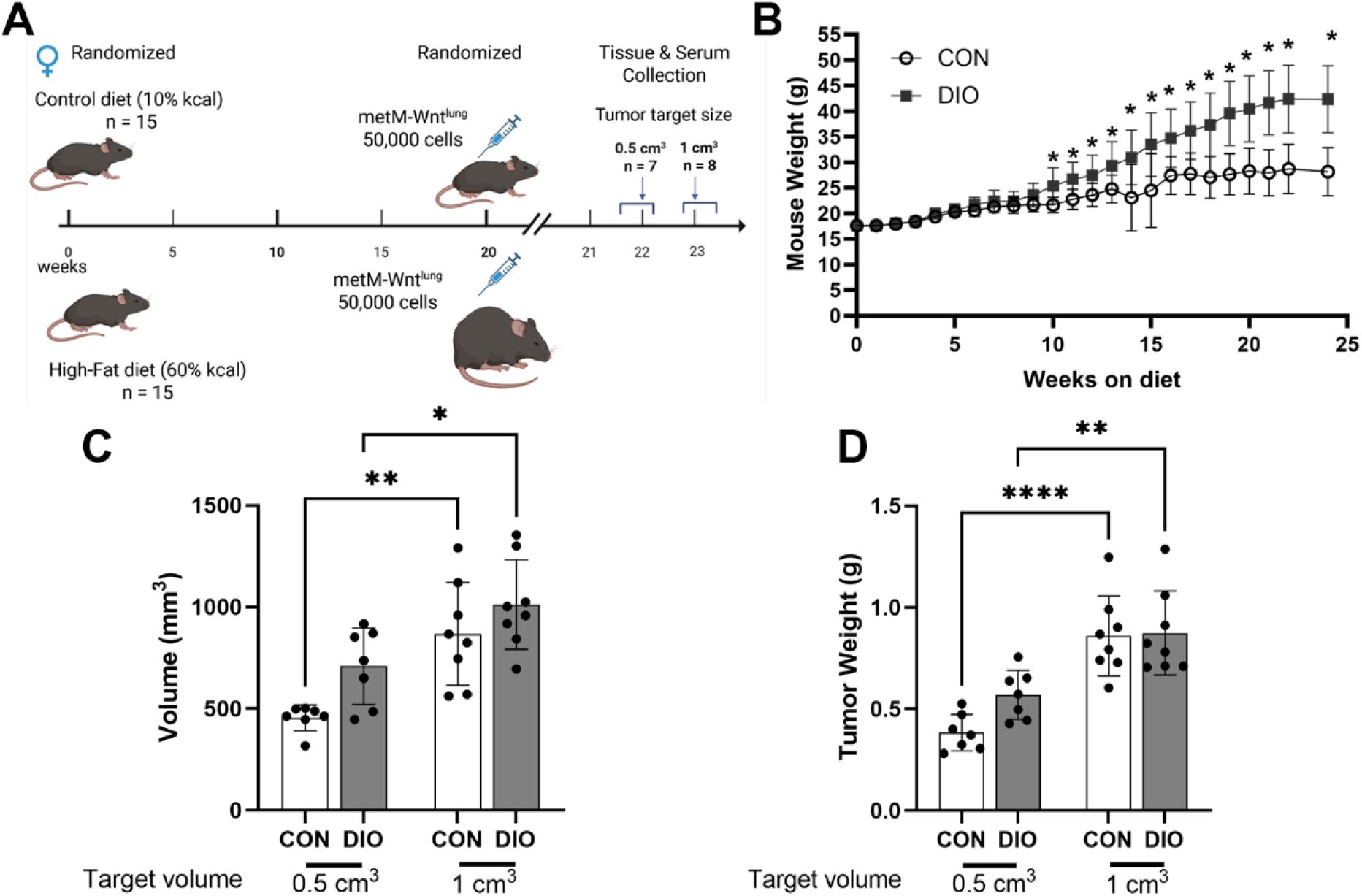
Experimental design and outcomes of orthotopic development of TNBC in female mice. **(A)** Experimental design. Female mice were fed either a control diet (CON; 10% kcal from fat) or a high-fat diet (DIO; 60% kcal from fat) for 20 weeks, inducing obesity in the DIO group. Subsequently, 50,000 metM-Wnt^lung^ cells were injected into the fourth mammary fat pad to induce mammary tumors. Tumors were collected when they reached ∼0.5 cm³ (“early”) or ∼1.0 cm³ (“late”) in volume. (B) Weekly body weights of CON and DIO mice throughout the study. (C) Tumor volume at collection in CON and DIO mice. (D) Tumor weight at collection in CON and DIO mice.

#### EV Isolation and Purification

From each mouse, EVs were purified separately from tumors (tumor-EVs) and VAT (VAT-EVs) following published guidelines (Welsh et al., 2024) and the method described by Crescitelli et al. (Crescitelli et al., 2021). Briefly, frozen tissues were weighed, placed on ice, and minced into ∼2 mm pieces. Tissue dissociation was performed in a ratio of 1 mL dissociation media per 0.2 g of tissue. For tumors, the dissociation media consisted of 2 mg/mL collagenase D and 80 U/mL DNase I in RPMI-1640; for VAT, the same enzymes concentrations were used in DMEM. Tissues were incubated in 15 mL tubes at 37 °C for 30 min on a nutating mixer (24 rpm), then filtered through 70 μm strainers. To maximize EV recovery, each tube was washed twice with 1 mL of the corresponding dissociation medium, and wash fractions were filtered and combined with the initial filtrate. After the washes, the combined filtrates were supplemented with protease inhibitors (1× final concentration) to stop enzymatic digestion. Samples were sequentially centrifuged at 300g for 10 min and 2,000g for 20 min at 4°C. The supernatant was transferred to polypropylene tubes (PN 4494, Selton Scientific) and ultracentrifuged at 16,500g for 20 min at 4°C in a Sorvall WX80 ultracentrifuge using a T-1270 fixed-angle titanium rotor (Cat# 08259, Thermo Fisher Scientific) with maximum acceleration and deceleration to pellet large EVs (LEVs) (Mogilevski et al.). The LEVs were resuspended in filtered PBS and stored at 4°C. The supernatant was then ultracentrifuged at 118,000g for 2.5 h at 4°C to collect the small EVs (SEVs), which were also resuspended in PBS and stored at 4°C.

To purify EVs and remove soluble contaminants, equal volumes of LEVs and SEVs from a given mouse and tissue were mixed and subjected to iodixanol gradient ultracentrifugation (OptiPrep, Cat. No. 1893, Serumwerk) with layers at 45%, 30%, and 10% (Crescitelli et al., 2021). After overlaying with 200 µL PBS, the gradient was centrifuged at 186,000g for 2.5 h at 4°C. The LEV/SEVs-rich band at the 30%/10% interface (at ∼ 1.058 -1.163 g/cm^3^) was collected for quality assessment and cell culture studies. The remaining EVs were stored at −80°C for proteomic analysis.

#### EVs Quality Assessment

##### Nanoparticle Analysis

The concentration and size of EVs from 6 mice in each group were evaluated using nanoparticle tracking analysis using a ZetaView® Z-NTA (Particle Metrix) instrument and ZetaView 8.04.02 software. The instrument was calibrated using 100 nm polystyrene nanostandard particles (Applied Microspheres) at a sensitivity level of 80 and minimum brightness of 20.

##### Western Blot

EVs were lysed in RIPA buffer with protease inhibitors, and protein concentration was determined by Bradford assay. A total of 20 µg of protein was separated by electrophoresis in a 4–12% SDS-PAGE gel, and transferred to PVDF membranes. After blocking with 5% milk in TBS-T, the membranes were incubated overnight at 4°C with primary antibodies against EV markers CD9 (Cat. No MA5-31980, Thermofisher), CD63 (Cat. No. SAB4301607, Sigma) or CD81 (Cat. No. MA5-33123, Thermofisher), and the non-EV markers calreticulin (Cat. No. 10292-1-AP, Proteintch) and calnexin (Cat. No mAb #2679, Cell signaling). Following 3 washes with TBS-0.5 % Tween-20, the membranes were incubated with fluorophore-conjugated secondary antibodies (LICORbio) and imaged using a LICOR Odyssey system.

##### Transmission Electron Microscopy

EVs were fixed in 2% glutaraldehyde in PBS for 30 min at room temperature. A 5–10 μL aliquot of the fixed EV suspension was placed onto a formvar/carbon-coated copper grid and allowed to adsorb for 20 min. Excess liquid was removed by blotting with filter paper. Grids were then washed in PBS and fixed with 1% glutaraldehyde for 5 min. After thorough washing with distilled water, samples were incubated with the contrast 2% uranyl acetate for 5–10 min. Grids were air-dried and imaged using transmission electron microscopy operated at 80–120 kV. EV morphology and size were assessed based on standard criteria (Chuo et al., 2018).

#### Proteomic Analysis

##### Sample Preparation

Purified EVs were processed for proteomic analysis in three separate batches of 12 unique samples each, based on specimen type (VAT, or tumor). A total of 45 µg of protein from each sample was dried via vacuum centrifugation, resuspended in 1X S-trap lysis buffer (5% SDS, 50mM TEAB), reduced with 5mM DTT at 56°C for 30 min, and alkylated with 15mM iodoacetamide for 45 min at room temperature. Samples were processed with S-trap spin columns (Protifi) following the manufacturer’s instructions then digested with trypsin (Promega). Resulting peptides were acidified to 0.5% trifluoroacetic acid and desalted using Thermo desalting spin columns. Eluates were dried via vacuum centrifugation, and peptide concentration was determined via Pierce Quantitative Fluorometric Assay. All samples were normalized to 0.25 µg/µL protein. A pooled sample was created by combining equal volumes of each unique sample within a batch to assess technical reproducibility.

##### LC/MS/MS and Proteomic Data Processing

Proteomes of the prepared EV samples were analyzed by LC-MS/MS using an Easy nLC 1200 system coupled to a Q-Exactive HF mass spectrometer (Thermo Scientific). Peptides were separated on an Easy Spray PepMap C18 column (75 μm × 25 cm, 2 μm particle size) over a 120-min gradient at a flow rate of 250 nL/min. The gradient consisted of a stepwise increase from 5% to 35%, and then to 45% mobile phase B (0.1% formic acid in acetonitrile), with mobile phase A being 0.1% formic acid in water. The Q-Exactive HF was operated in data-dependent acquisition mode, selecting the 15 most intense precursor ions for HCD fragmentation. Full MS scans (m/z 300–1600) were acquired at 120,000 resolution, with an AGC target of 3 × 10 ions and a maximum injection time of 100 ms. MS/MS scans were acquired at 15,000 resolution with an AGC target of 1 × 10 ions, maximum injection time of 75 ms, a normalized collision energy of 27%, and an isolation window of 1.6 m/z. Peptide match was set to "preferred," and precursors with unknown charge states, or charges of 1 or ≥8, were excluded from fragmentation. Unique samples were analyzed in random order, and the pooled sample was analyzed intermittently to assess instrument variability.

Raw data were analyzed using SequestHT within Proteome Discoverer v2.5 (Thermo Scientific). Spectra were searched against the reviewed mouse Uniprot proteome database (UP000000589, downloaded July 2021, 17,051 sequences) appended with a common contaminants database (MaxQuant; 245 proteins). Samples were searched as three separate batches based on specimen source (VAT, or tumor), including their respective pooled-sample replicates. Search parameters included trypsin specificity allowing up to two missed cleavages, fixed carbamidomethylation of cysteine, and variable modifications of methionine oxidation and N-terminal acetylation. Peptide and protein identifications were filtered at a 1% and 5% false discovery rate (FDR), respectively. Proteins identified with fewer than two unique peptides and/or with more than 50% missing values across samples were excluded. Protein-level label-free quantification (LFQ) intensity values were log₂-transformed, and sample-specific abundance counts were used to estimate peptide-level evidence. The minimum abundance count per protein across all samples, plus a pseudocount of one, was used to model intensity-dependent variance.

Differential protein expression analysis was performed using the DEqMS package (Zhu et al., 2020) in R. Proteins with an adjusted p-value (FDR) < 0.05 were considered significantly differentially expressed. Principal component analysis (PCA) and hierarchical clustering were performed to evaluate sample relationships and expression patterns.

Differentially expressed proteins were compared against the ExoCarta database (www.exocarta.org) by matching gene symbols or UniProt IDs to identify proteins associated with EVs. Overlap with ExoCarta entries was quantified. Gene Ontology (GO) enrichment analysis was performed using DAVID in R, focusing on Biological Process (BP), Cellular Component (CC), and Molecular Function (MF) categories. Enrichment significance was determined using a Benjamini–Hochberg adjusted p-value < 0.05. Enriched pathways were visualized using dot plots, where dot size reflected the number of genes per term and dot color represented the adjusted p-value.

#### Isolation and Activation of CD8**D** T Cell from Murine Spleen

CD8 T cells were isolated from pooled spleens of 7 age-matched (30 weeks old) C57BL/6 lean females mice, using magnetic negative selection (STEMCELL Technologies, Vancouver, BC, Canada, Cat. No 19853) according to the manufacturer’s protocol as previously described (Kiernan et al., 2024). Isolated cells were activated with 1 μg/mL anti-CD3 (BioLegend, San Diego, CA, Cat No 100340) and 5 μg/mL anti-CD28 (BioLegend, Cat No 102116) antibodies for 48 h in 96-well flat bottom plates (200,000 cells/well, 200µL total).

#### Confocal Imaging of EV Uptake in CD8^+^ T or Mammary Cancer Cell Lines

Isolated CD8 T cells were activated in ibidi chambers (Ibidi, Cat.No 81817, final volume: 100 µL) coated with 1 μg/mL anti-CD3 (BioLegend, San Diego, CA, Cat No 100340) and 5 μg/mL anti-CD28 (BioLegend, Cat No 102116) antibodies for 48 h prior treatment with EVs.

E-Wnt and M-Wnt cells were used as EV recipient lines to determine whether EV uptake and metabolic responses depend on the phenotypic state of the tumor cell (epithelial-like versus mesenchymal-like) rather than representing a uniform tumor response. E-Wnt and M-Wnt cells were cultured on glass coverslips in 24-well plates (final volume: 200 µL) in complete RPMI 1640 medium supplemented with 10% FBS, 1% HEPES, 1% penicillin/streptomycin, and 1% glutamine. A total of 50 µg of EVs or 0.02 μm-filtered PBS (negative control) were stained with 10 μM of the red fluourescent lipophilic dye that incorporates into lipidic membranes PKH26 (Mini26, Sigma) for 10 min at 37 °C, followed by ultracentrifugation at 100,000 × g for 1 h to remove unbound dye.

Stained EVs were resuspended in 150 µL of filtered PBS and added to CD8 T cells, E-Wnt cells, or M-Wnt cells at 1:4 ratio (EV:cell culture volume) and co-cultured for 24 h. After co-culture, the medium was removed, and cells were gently rinsed with warm PBS. The cells nuclei and plasma membranes were stained with Hoechst 33342 and 2× CellBrite Fix 640 (Biotium, Cat. No. 30089) for 15 min at 37 °C, respectively. Cells were then gently washed twice with PBS and fixed with 2% paraformaldehyde for 10 min, followed by two washes with PBS. Coverslips were mounted with ProLong gold antifade mounting medium (Invitrogen, Cat. No. P36930) and allowed to dry. T cells were mounted with ibidi mounting media (Ibidi, Cat. No 50001). Imaging was performed on a Zeiss 810 confocal microscope, and image processing was conducted using Fiji (Schindelin et al., 2012).

#### CD8□ T Cell Functional Assay by Flow Cytometry

During activation (48 h), CD8 T cells were treated with 50 μg/mL tumor- or VAT-EVs isolated from early- or late-stage mammary tumors. Unstimulated CD8 T cells were treated in parallel to assess EV effects independent of activation. Cells were then harvested and stained as previously described (Kiernan et al., 2023) using Cytofix/Cytoperm kit (BD Biosciences, San Jose, CA) following the manufacturer’s protocol. The following fluorochrome-conjugated antibodies were used: CD8-PE (BioLegend, Cat. No 100708, 1:400), CD44 PE/Cy7 (BioLegend, Cat. No 103030, 1:400), granzyme B-APC (BioLegend, Cat. No 372204, 1:200), IFN-γ-PE/Cy7 (BioLegend, Cat. No 505826, 1:200), and TNF-AF647 (BioLegend, Cat. No 506314, 1:200). Flow cytometry was performed on an Accuri C6 flow cytometer (BD Biosciences), and data were analyzed using FlowJo software (BD Biosciences). The geometric mean fluorescence intensity (gMFI) of each marker was quantified to assess changes in CD8 T cell activation and effector molecule expression.

#### Seahorse Metabolic Flux Assay in CD8**D** T Cells and Mamary Cancer Cell Lines

Activated CD8 T cells were treated with 50 µg/mL of purified EVs in a 24-well plate (1 million cells/well, 1 mL total volume) and assessed for metabolic function using the Seahorse XFe96 Analyzer (Agilent). Prior to the assay, CD8 T cells were washed and resuspended in Seahorse XF RPMI 1640 medium (Agilent, Santa Clara, CA), then plated at a density of 250,000 cells per well (50 µL) in Seahorse XFe96 cell culture microplates pre-coated with Cell-Tak (Corning, Corning, NY). Plates were centrifuged at 200 × g for 1 min and incubated for 30 min in a humidified 37 °C incubator without CO . Subsequently, 130 µL of Seahorse XF RPMI 1640 medium was added to each well, followed by an additional 20-min incubation under the same conditions.

E-Wnt and M-Wnt cells were seeded onto Seahorse XFe96 plates 96 h prior to the assay at densities of 15,000 and 20,000 cells per well, respectively, in complete growth medium (CM). The following day, cells were washed three times with PBS, and the medium was replaced with EV-depleted medium, prepared with ultracentrifuged FBS (UCF) at 100,000 × g for 18 h. Cells were then treated with 25 µg of EVs and co-cultured for 48 h. On the day of the Seahorse assay, cells were washed and incubated in Seahorse XF RPMI 1640 medium for 1 h at 37 °C in a non-CO incubator to allow for temperature and pH equilibration.

Oxygen consumption rate (Filippone et al.) and extracellular acidification rate (ECAR) were measured under basal conditions and following the sequential injection of the following compounds (each purchased from Sigma-Aldrich): 1 µM oligomycin, 0.5 µM (or 1 µM) fluoro-carbonyl cyanide phenylhydrazone (FCCP), and a combination of 0.75 µM (or 1 µM) rotenone with 1.5 µM (or 1 µM) antimycin A (parenthetical concentrations were used for cancer cells). Data were analyzed using Wave software (Agilent) to evaluate mitochondrial respiration.

#### Statistical Analysis

Statistical analyses were performed using GraphPad Prism v10.4.2 and R v4.4.2. Outlier detection was conducted using Dixon’s Q test, but no data points were excluded. Data are presented as mean ± SEM. Group differences were assessed using one-way or two-way analysis of variance (ANOVA), as indicated. PCA was performed and visualized using the faxtoextra package in R (Kassambara and Mundt, 2020). A p-value of <0.05 was considered statistically significant.

## Results

### Animal model of obesity and orthotopic TNBC development

We used a well-described mouse model of obesity and TNBC in which control and high-fat diet results in lean (CON) and obese (DIO) phenotypes, respectively, and orthotopically transplanted murine TNBC cells (metM-Wnt^lung^) induce mammary tumors (Zheng et al., 2013, O’Flanagan et al., 2017) (Figure 1A). In our study, after 20 weeks on diet, the average body weights were 28.3 g for CON mice and 40.5 g for DIO mice (Figure 1B). Following orthotopic transplantation of metM-Wnt^lung^ cells, CON and DIO mice were randomized to allow mammary tumors to grow. Tumors and VAT were harvested when tumors reached approximately 0.5 cm^3^ (classified as “early tumors”; n=7) or 1.0 cm^3^ (classified as “late tumors”; n=8). Upon tumor harvest, we detected no significant difference in tumor volume (Figure 1C) or tumor weight (Figure 1D) between dietary groups for early (i.e. CON-0.5 vs DIO-0.5) or late tumors (i.e. CON-1 vs DIO-1). In contrast, within each dietary group, tumor volume and weight significantly differed between early and late tumors (e.g. CON-0.5 vs CON-1 or DIO-0.5 vs DIO-1, Figure 1C, D). Together, this indicated consistency and minimal variability during tumor measurements and sample collection.

### Isolation and characterization of EVs

EVs were isolated and purified from tumors and VAT (tumor-EVs and VAT-EVs; Figure 2A) by ultracentrifugation in combination with an Iodixanol gradient. By transmission electron microscopy (Figure 2B), EVs appeared as round, membrane-bound structures ranging from 50 to 200 nm in diameter and displaying a characteristic cup-shaped or spherical morphology that has been reported to result from dehydration and staining artifact during sample preparation (Chuo et al., 2018). In addition, Tetraspanin EV markers (CD9 and CD63 or CD81) were detected by western blot (Figure 2C) across multiple fractions collected during EV isolation, including pellets obtained after each centrifugation step and the LEV, SEV, and combined LEV/SEV fractions (following flotation gradient separation) derived from VAT and tumor tissues of CON and DIO mice bearing early (CON-0.5, DIO-0.5) or late (CON-1, DIO-1) tumors. In western blots performed to verify EV purity, calnexin and calreticulin were undetectable in EVs from CON-0.5 and DIO-0.5 mice bearing early tumors and faintly detected in EVs preparations from mice bearing late tumors (CON-1 and DIO-1), but calreticulin was more abundant in tumor-EVs from DIO-1 mice (Supplemental Figure 1). By nanoparticle tracking analysis, average tumor-EV diameters (Figure 2D) were 143.1 nm for CON-0.5, 134.3 nm for DIO-0.5, 127.5 nm for CON-1, and 120 nm for DIO-1, with a significant difference observed between tumor-EVs from CON-0.5 and DIO-1 groups. In contrast, VAT-EV diameters were comparable across diet groups and tumor stages, with mean sizes of 134.6 nm (CON-0.5), 131.2 nm (DIO-0.5), 134.4 nm (CON-1), and 133.0 nm (DIO-1) (Supplementary Table 1). Within the EV enriched fraction isolated at the 30%/10% Iodixanol interface (∼1.058–1.163 g/cm³), tumor-EVs exhibited a shift toward smaller average diameters across groups, whereas VAT- EV size distributions remained largely stable.

**Figure 2.**
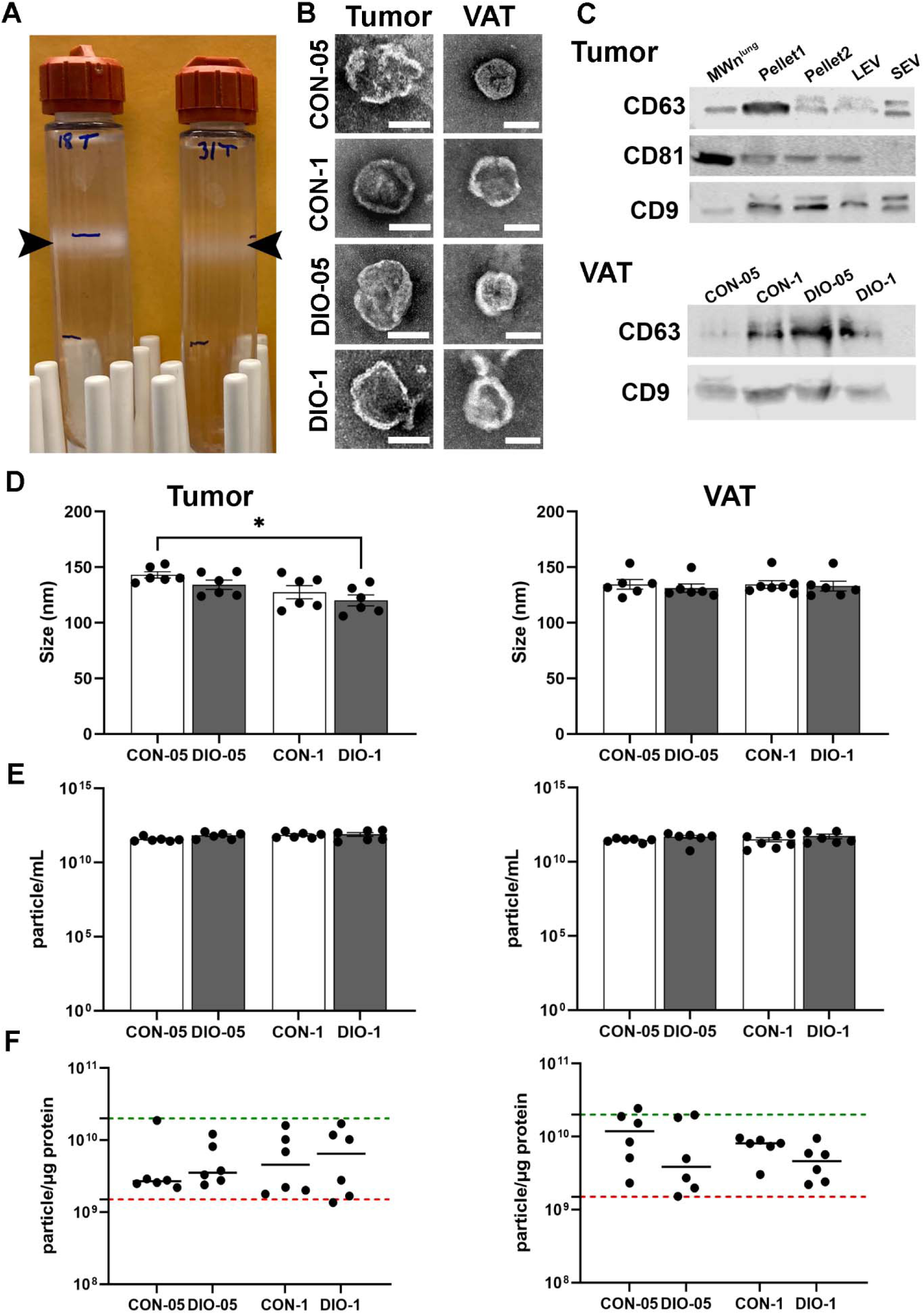
Purification and characterization of EVs. **(A)** Representative flotation gradient following ultracentrifugation. The white band at the 30–10% interface (1.058 -1.163 g/cm^3^), containing the collected EV fraction, is indicated by arrowheads. **(B)** Representative transmission electron micrographs of EVs isolated from tumor and VAT. Scale bar = 100 nm (**C)** Western blot analysis showing the presence of EV markers CD9 and CD63 or CD81 in EVs, as indicated in each panel. Protein lysate from met-M-Wnt^lung^ (MMWnt^lung^) cells was used as a positive control. **(D)** Size and **(E)** concentration of EVs isolated from tumor and VAT of lean (CON) or diet-induced obesity (DIO) mice bearing early (0.5 cm³) or late (1.0 cm³) tumor. Data represent mean ± SEM, measured using the ZetaSizer Nano ZS (n = 6). **(F)** Particle-to-protein ratio (particles/µg protein) as a measure of EV preparation purity for each condition (n = 6). Dashed lines indicate the acceptable range for pure EVs (2×10_ to 2×10¹_ particles/µg protein). Each symbol represents an individual sample. Data are shown as mean ± SEM. Asterisks indicate significant differences determined by one-way ANOVA with multiple comparisons (*p < 0.05, **p < 0.01, ***p < 0.001).

To assess the purity of the EV preparations, we used the EV concentrations (particle/mL, by nanoparticle tracking analyses; Figure 2E; Supplementary Table 1) and the total protein concentration to calculate the particle-to-protein ratio (Figure 2F). Ratios of 2×10 to 2×10¹ particles/mL/µg protein are considered indicative of pure EV preparations (Webber and Clayton, 2013, Bellotti et al., 2021). The ratios for tumor- and VAT-EVs fell within this range, consistent with acceptable purity.

### Global Proteomic Analysis of Tumor-EVs and VAT-EVs

To determine how obesity and tumor development influence the proteomic composition of EVs, tumor- and VAT-EVs were individually analyzed by mass spectrometry (Figure 3A).

**Figure 3:**
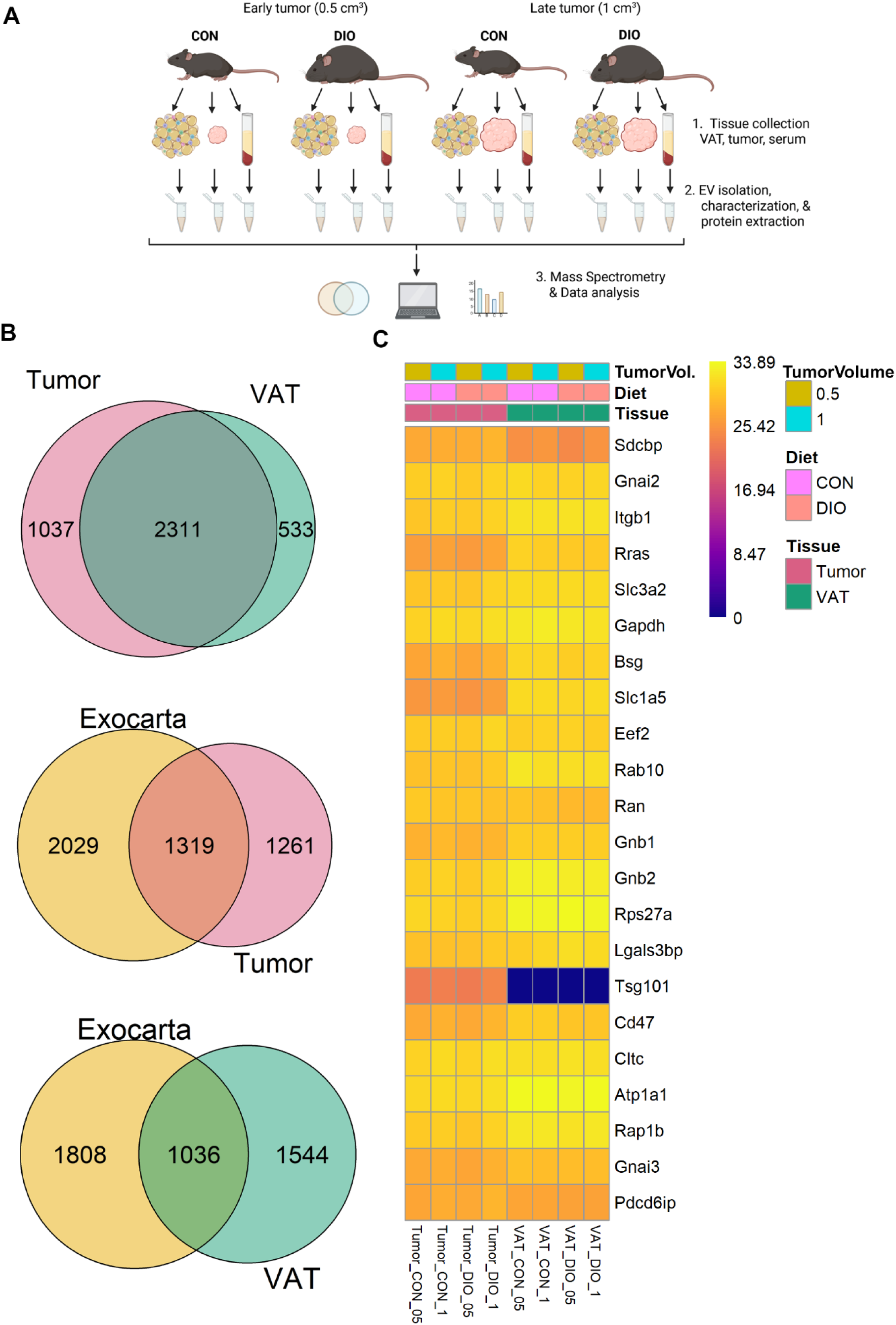
Global EV proteomic Analysis. **(A)** Schematic representation of the experimental workflow. Created in BioRender (https://BioRender.com/0tec6oe). Female mice fed either a control diet (CON) or a diet-induced obesity (DIO) diet and bearing mammary tumors were sacrificed when tumor volumes reached 0.5 cm³ (early tumors) or 1 cm³ (late tumors). Tumors and VAT were collected for EV isolation and subsequent proteomic analysis using mass spectrometry. **(B)** Venn diagrams showing proteins identified in EV samples compared across tumor and VAT as well as proteins annotated in the ExoCarta database. **(C)** Heatmap displaying the relative abundance of EV markers proposed by Kugeratski et al.

Global proteomic profiling identified 3,348 proteins in tumor-derived EVs and 2,844 proteins in VAT-derived EVs (Figure 3B). Tumor- and VAT-EVs shared 2,071 proteins, representing 53% of the combined EV proteome (Figure 3B, top Venn diagram), indicating substantial overlap between the two EV sources.

We next cross-referenced the global EV proteomic datasets with ExoCarta using a curated mouse EV protein set comprising 2,580 unique entries after removal of duplicate annotations (Figure 3B). A substantial fraction of tumor- and VAT-derived EV proteins overlapped with ExoCarta annotations, consistent with enrichment of canonical EV-associated proteins, while additional proteins detected in our datasets were not represented in the database.

To validate the EV-proteomic data, we screened our results for the presence of 22 proteins described as the proteomic signature for EVs. These proteins have previously been identified as highly abundant in EVs derived from a variety of cellular compartments and isolated using diverse procedures (Kugeratski et al., 2021). To simplify the analysis, we averaged the relative abundance (log2-LFQ) of the samples from the same dietary group and tumor size, then used an R script to develop a heat map (Figure 3C). Tumor- and VAT-EVs demonstrated a high degree of overlap with the 22 known EV markers (specifically, demonstrating 22 and 21 of the markers, respectively), consistent with both EV populations being representative of bona fide, purified EVs suitable for downstream proteomic analyses. The one known protein marker that VAT-EVs lacked was TSG101 (Figure 3C). Kugeratski et al. also described a list of 1,285 genes corresponding to 1,212 proteins commonly found in EVs (Kugeratski et al., 2021). The global protein lists for tumor-, VAT-EVs in our study contained 72% and 58% of these genes, respectively (Supplementary Figure 2).

### Distinct Effects of Tumor Progression and DIO on the Proteomic Landscape of Tumor-and VAT-EVs

PCA was used to evaluate the relative contributions of tumor progression and DIO to the proteomic composition of tumor- and VAT-EVs. In tumor-derived EVs, principal component 1 accounted for 46% of the variance (P = 0.003) and clearly separated EVs isolated from early (0.5 cm³) versus late (1.0 cm³) tumors (Figure 4A), indicating pronounced proteomic remodeling of tumor EVs with tumor progression. In contrast, DIO versus control lean status did not significantly influence the tumor EV proteome (P = 0.449; Figure 4B).

**Figure 4.**
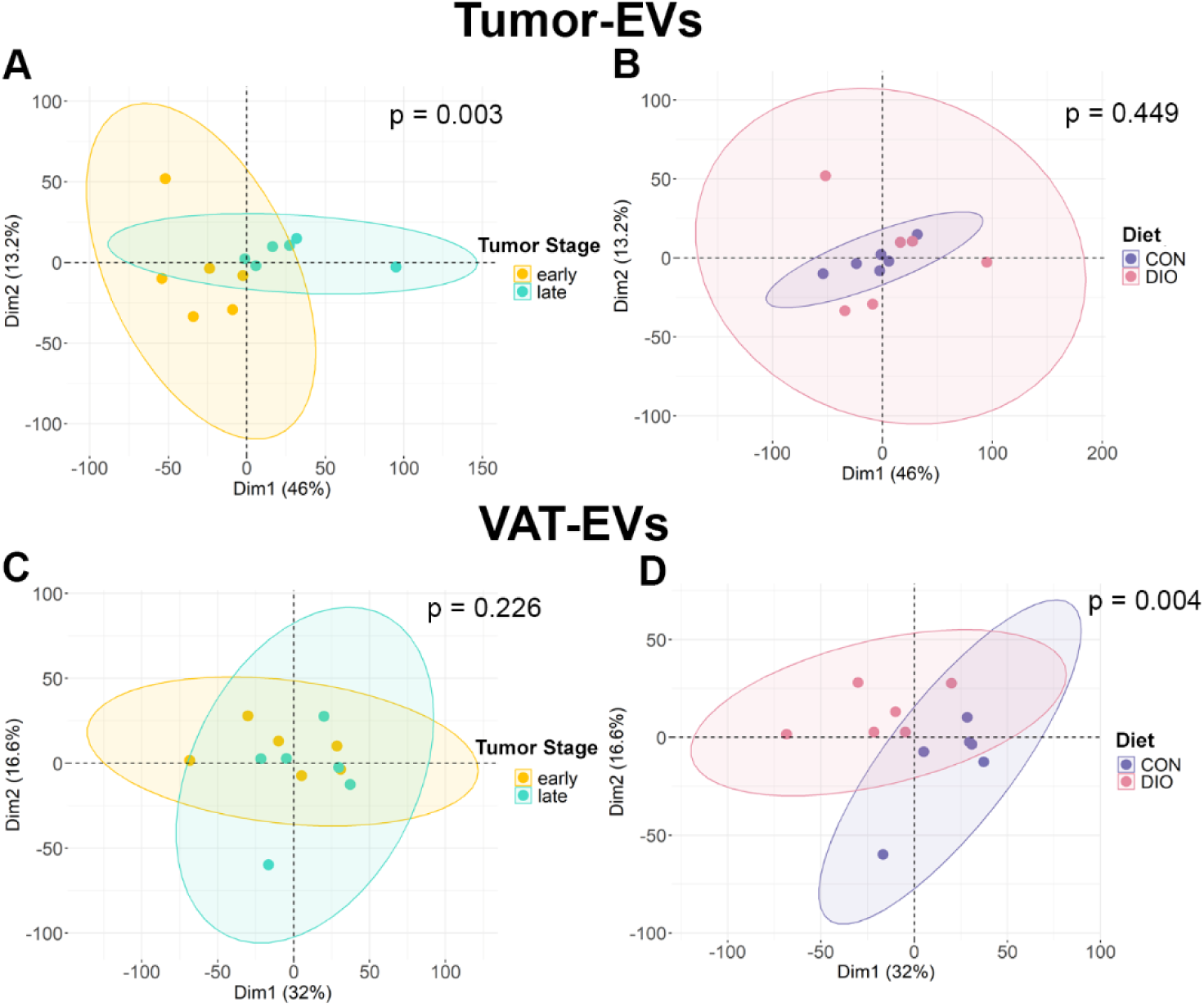
Principal component analysis (PCA) of EV proteomes from tumor and VAT reveals distinct effects of tumor size and diet. (A, **B)** PCA of tumor-derived EV proteomes. **(A)** Component 1 accounted for 46% of the total variance and significantly separated EVs from early (0.5 cm³) and late (1.0 cm³) tumors (P = 0.003), indicating tumor progression strongly alters EV protein composition. **(B)** No significant clustering was observed based on diet in tumor-derived EVs (P = 0.449). **(C, D)** PCA of VAT-derived EV proteomes. In contrast to tumor EVs, **(C)** tumor stage has not major impact on VAT-EVs proteomic composition. **(D)** In VAT-EVs, diet was the primary factor driving differences. Clear separation was observed between EVs from control diet (CON) and diet-induced obesity (DIO) groups, suggesting that systemic metabolic status shapes VAT-EV composition. n = 6 in each group, p-values indicated in the corresponding panel.

In VAT- EVs, DIO status was the primary determinant of proteomic variation (P = 0.004; Figure 4D), whereas tumor size had no significant effect (P = 0.226; Figure 4C). Together, these analyses demonstrate that tumor progression predominantly shapes the proteomic landscape of tumor- EVs, whereas diet-induced obesity is the dominant driver of proteomic changes in VAT-derived EVs.

### Differential Protein Expression in EVs During Tumor Progression in Lean and DIO Mice

To identify changes in the proteomic profile of EVs during tumor progression, we conducted differential protein expression analysis using DEqMS—a robust statistical method optimized for mass spectrometry data (Zhu et al., 2020). Pairwise comparisons were performed within each diet group for tumor- and VAT-EVs from mice bearing early (0.5 cm³) or late (1 cm³) tumor (i.e., CON-1 vs. CON-0.5 and DIO-1 vs. DIO-0.5). Volcano plots (Figure 5) were generated to highlight EV proteins exhibiting at least a log₂ fold change ≥ +1.3 (2.5-fold change) with a P-value ≤ 0.01 within each group.

**Figure 5.**
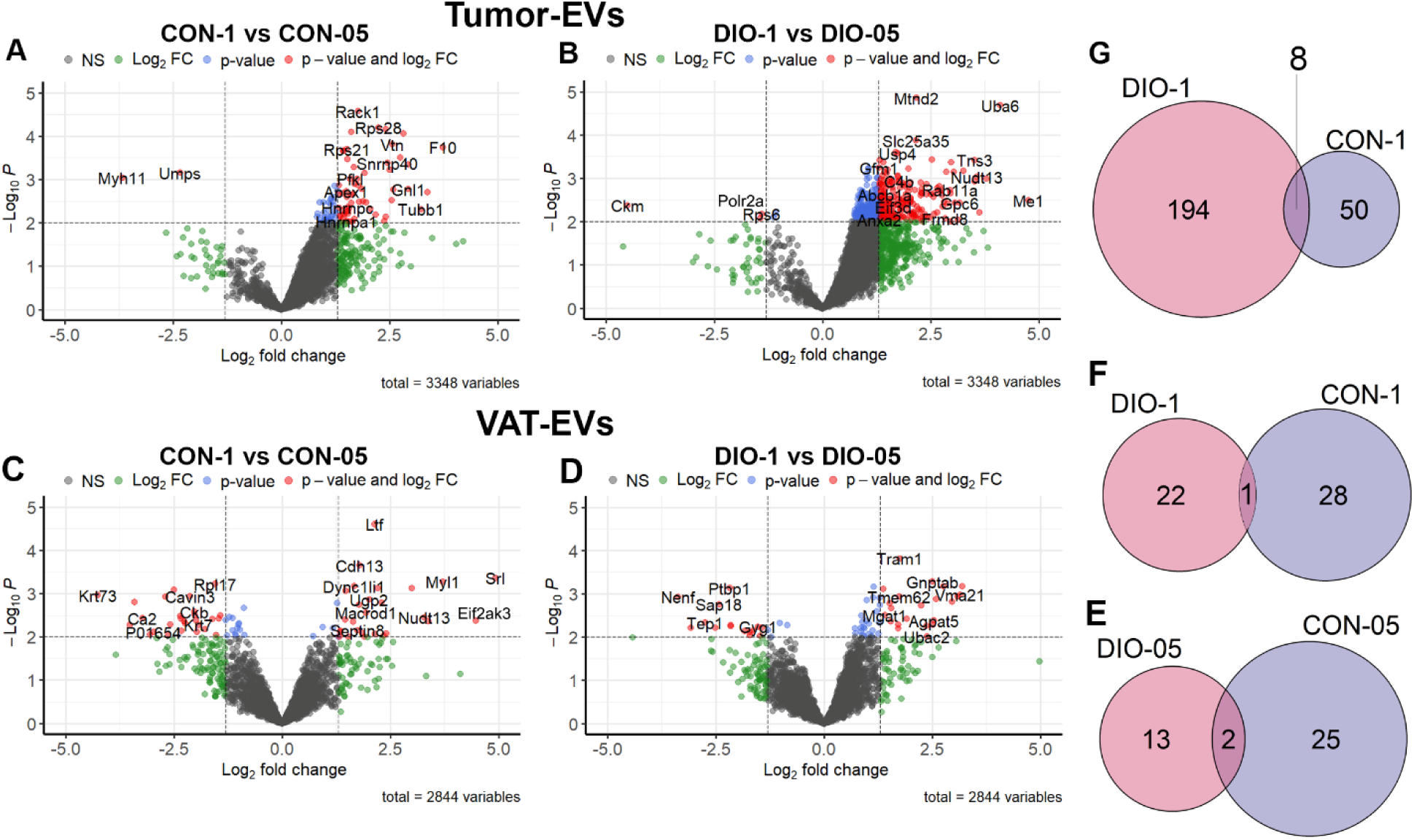
Differential Protein Expression in EVs During Tumor Progression Within Dietary Groups. (A, **B)** Volcano plots showing differentially expressed proteins in tumor-derived EVs isolated at early (0.5 cm³) and late (1.0 cm³) tumor stages in control (CON) and diet-induced obesity (DIO) mice. **(C, D)** Volcano plots of VAT-derived EVs from CON and DIO mice at early and late tumor stages. Proteins with log₂ fold change ≥ +1.3 and P ≤ 0.01 are highlighted in red. Distinct proteomic signatures illustrate diet-and stage-dependent alterations in EV composition. **(E–G)** Venn diagrams showing overlap among upregulated proteins across conditions. **(E)** VAT-derived EVs from DIO-05 and CON-05 share Mettl9 and P01654. **(F)** VAT-derived EVs from DIO-1 and CON-1 share upregulation of Eif2ak3. **(G)** Tumor-derived EVs from DIO-1 and CON-1 share a core set of eight upregulated proteins (Itih4, Itga2b, Procr, Hspb1, Gpld1, Hspa1a, Adprh, Enpp1).

In early tumors, tumor-EVs from CON-0.5 showed upregulation of proteins involved in cytoskeletal organization and nucleotide metabolism— MYH11 (myosin heavy chain 11) and UMPS (uridine monophosphate synthetase) (Figure 5A). In contrast, tumor-EVs from DIO mice (DIO-0.5, Figure 5B) were enriched in proteins associated with energy metabolism, non-coding RNA processing and protein synthesis, including CKM (creatine kinase M-type), POLR2A (RNA polymerase II subunit A) and RPS6 (ribosomal protein S6, a key component of mTOR signaling), as summarized in Supplementary Table 2.

In late tumors, tumor-EVs from CON-1 exhibited marked upregulation of proteins related to coagulation (e.g., factor 10 and prothrombin), extracellular matrix (ECM) remodeling (e.g., Timp2), metabolic adaptation (e.g., PFKL), and stress response (e.g., DNAJC5) (Figure 5A, Table 1 and Supplementary Table 2). Conversely, tumor-EVs from DIO-1 mice demonstrated significant increase in proteins associated with lipid metabolism (e.g., MGII, LIPE), immune modulation (e.g., Tlr7), metabolic reprogramming (e.g., CYP2d11), and ECM remodeling (Figure 5B, Table 1 and Supplementary Table 2).

By comparison, VAT-EVs from control lean mice bearing early tumors (CON-0.5) showed upregulation of proteins involved in immune signaling, structural integrity, and metabolic regulation, including LRPAP1, RPS13, MSLN, RPL17, and TMEM214 (Figure 5C). In contrast, VAT-EVs from DIO mice with early tumors (DIO-0.5) were enriched in proteins such as KARS1, PZP, GYG1, PARP4, and MYP, consistent with obesity-associated alterations in extracellular matrix remodeling, metabolic adaptation, and intracellular signaling (Table 2 and Supplementary Table 3).

In control mice (CON-05) bearing early tumors, VAT-EVs showed upregulation of proteins involved in immune signaling, structural integrity, and metabolic regulation—such as LRPAP1, RPS13, MSLN, RPL17, and TMEM214 (Figure 5C). In contrast, VAT-EVs from the DIO-0.5 group were enriched in proteins like KARS1, PZP, GYG1, PARP4, and MYP, consistent with obesity-induced changes in ECM remodeling, metabolic adaptation, and intracellular signaling. (Table 2 and Supplementary Table 3).

At the late tumor stage, the VAT-EVs from CON-1 mice were enriched in proteins linked to cytoskeletal and contractile functions—including TPM12, TNNT3, SRL, EIF2AK3, and MYL1—which may reflect enhanced cellular dynamics and tissue remodeling in response to tumor progression (Figure 5D, Table 2 and Supplementary Table 3). Conversely, VAT-EVs from DIO-1 mice exhibited increased protein levels of CAPN5, VMA21, YIPF3, and SLC38a7, indicative of altered stress responses and metabolic regulation under obese conditions (Supplementary Table 3).

In addition to these distinct proteomic profiles, our analysis identified overlapping protein signatures across EV sources and diet groups (Table 3). During early tumor development, VAT-derived EVs from both CON-0.5 and DIO-0.5 mice consistently upregulated METTL9 and P01654, suggesting that certain tumor-associated alterations in VAT-EV cargo are conserved regardless obesity state (Figure 5E). At the late tumor stage, VAT-EVs from both groups (CON-1 and DIO-1) commonly upregulated EIF2AK3, consistent with a shared stress response or metabolic adaptation (Figure 5F).

Similarly, at the late tumor stage, tumor-derived EVs from both control lean and DIO mice (CON-1 and DIO-1) shared a core set of eight upregulated proteins (Figure 5G)—ITIH4, ITGA2B, PROCR, HSPB1, GPLD1, HSPA1A, ADPRH, and ENPP1—implicated in ECM remodeling, immune modulation, and metabolic reprogramming. Notably, AGPAT5 was the only protein consistently upregulated in both VAT- and tumor-derived EVs from DIO mice at the late tumor stage, highlighting a potential role for lipid metabolism in shaping a protumor microenvironment under obese conditions (Table 3).

Collectively, these results indicate that EV protein composition evolves with tumor progression and is differentially influenced by obesity. The overlapping and distinct proteomic signatures suggest the existence of a conserved EV-mediated signaling framework that coordinates metabolic and immune responses during tumor progression, with obesity selectively amplifying lipid-associated pathways.

### Obesity Regulation of EV Protein Profiles

To investigate how obesity influences the proteome of tumor- and VAT-EVs during tumor development, we compared the proteomic profile of EVs purified from lean and obese mice bearing early (DIO-0.5 vs. CON-0.5) and late (DIO-1 vs. CON-1) tumors. In early tumor-EVs, we identified 22 upregulated and 14 downregulated proteins in the DIO-0.5 vs. CON-0.5 comparison (Figure 6A). These changes in EV protein cargo suggest that obesity-driven tumors exhibit increased aggressiveness, altered immune interactions and surveillance, impaired metabolic flexibility, and mitochondrial dysfunction, as further supported by Gene Ontology (GO) pathway analysis (Figure 6A). In late tumor-EVs, 59 proteins were upregulated in DIO-1 vs. CON-1 comparison, with no proteins significantly downregulated (Figure 6B). Many of these proteins—including MGLL, LIPE, CD36, FABP4, and ACSL1—are key regulators of lipid uptake, activation, and catabolism, while AGPAT2, AGPAT5, CHPT1, and GPD1 support phospholipid biosynthesis. Additional proteins such as COX6A1, MAOB, ABCD2, ALDH6A1 and PC are linked to mitochondrial metabolism and redox balance, and matrix remodeling is reflected by upregulation of COL1A1, COL5A2, and PRELP. Other identified upregulated proteins, including ADIPOQ, DPT, and SOD3, suggest broader effects influencing systemic metabolism, tissue remodeling and redox status, respectively. Upregulated proteins including SLC27A1, PC, MTOR, and MACF1 further imply enhanced tumor cell proliferation, metabolic flexibility, and invasive potential, consistent with a more aggressive tumor phenotype.

**Figure 6:**
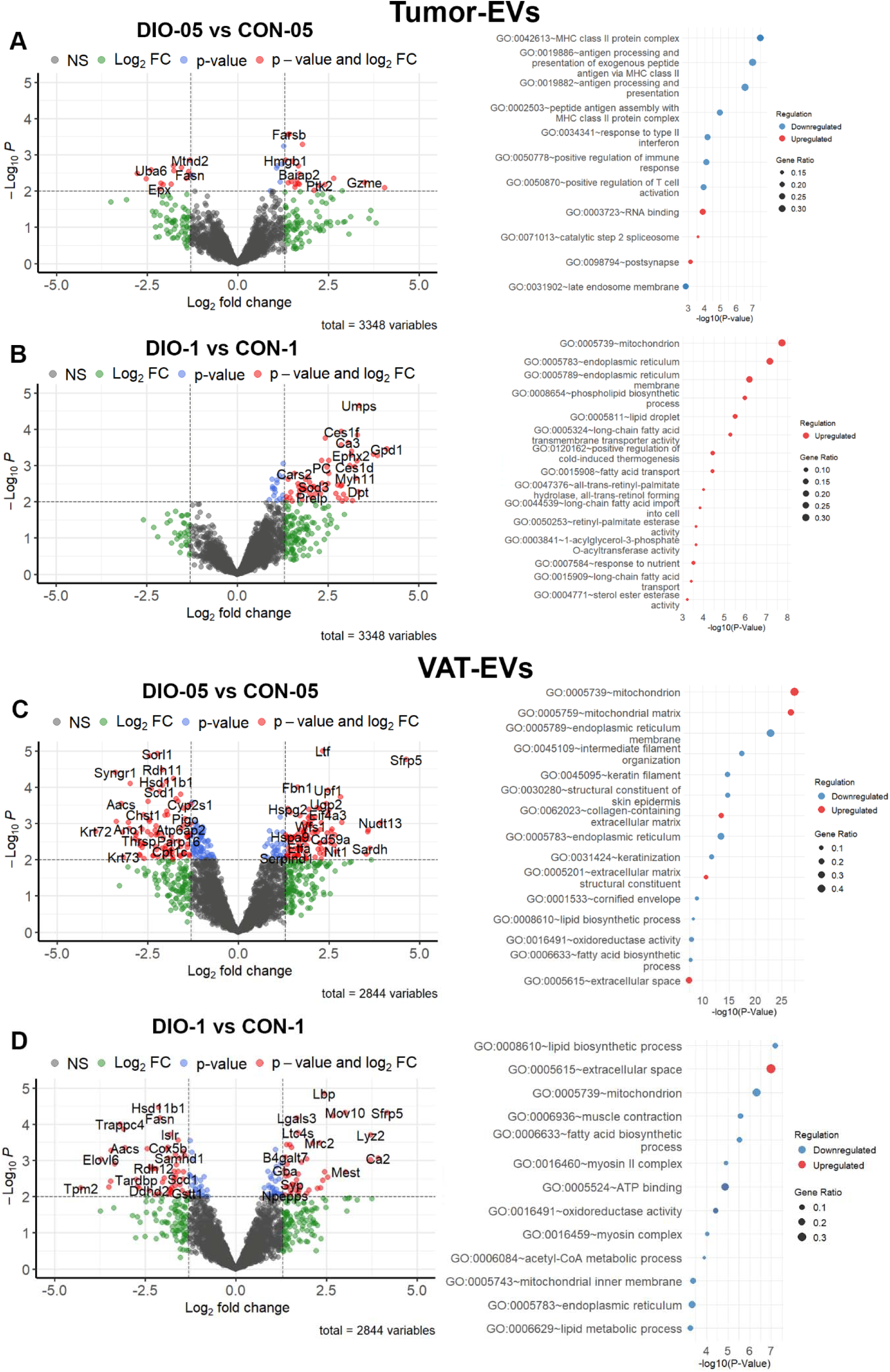
Figure 2. Diet-Dependent Differences in Tumor- and VAT-Derived EV Proteomes at Equivalent Tumor Stage. (A, **B)** Differential expression analysis of tumor-derived EVs from diet-induced obesity (DIO) and control (CON) mice at (A) early (0.5_cm³) and (B) late (1.0_cm³) tumor stages. (C, D) Differential expression analysis of VAT-derived EVs from DIO and CON mice at early (C) and late (D) tumor stages. Each panel includes a volcano plot of differentially expressed proteins (log₂ fold change ≥_+1.3; P ≤_0.01, highlighted in red) and a corresponding gene ontology (GO) enrichment analysis showing the top 15 upregulated (red) and downregulated (blue) pathways. The dot size represents gene ratio.

Consistent with these proteomic changes, GO analysis of the upregulated proteins in late tumor-EVs, revealed enrichment of pathways involved in phospholipid biosynthesis, long-chain fatty acid transport, nutrient sensing, and cold-induced thermogenesis. Functional categories included lipid hydrolase, acyltransferase, and oxidoreductase activity, while enriched cellular components—such as the endoplasmic reticulum, mitochondria, lipid droplets, and extracellular space—support a metabolic role of the EVs. These findings suggest that obesity reprograms tumor-EV composition to promote lipid metabolism, mitochondrial adaptation, and stromal remodeling.

The analysis of VAT-EVs purified from mice bearing early tumors identified 103 upregulated and 94 downregulated proteins in the DIO-05 vs. CON-05 comparison (Figure 6C). GO analysis of VAT-EVs showed upregulation of multiple components involved in extracellular matrix remodeling, mitochondrial pathways, the tricarboxylic acid (TCA) cycle, and fatty acid oxidation These findings suggest that VAT-EVs from DIO-0.5 mice could be more metabolically active. In the VAT-EVs from mice bearing late tumors, we identified 43 upregulated and 55 downregulated proteins (Figure 6D). GO analysis showed upregulation of extracellular space (GO:0005615) and extracellular region (GO:0005576), possibly indicating that VAT-EVs are more actively secreted into circulation in late-stage tumors and carry more cytoplasmic-derived molecules (GO:0005829). These VAT-EVs also exhibited upregulation of ATP binding and oxidoreductase activity proteins, reflecting altered energy metabolism and stress response in VAT during late tumor development. However, unlike VAT-EVs from mice bearing early tumors, where proteins involved in mitochondrial metabolism were upregulated, VAT-EVs from mice bearing late tumor showed downregulation of mitochondrial pathways (GO:0005739, GO:0005743) and the mitochondrial inner membrane (GO:0005743), indicating a metabolic shift in VAT-EVs. In addition, lipid metabolism and structural protein content were downregulated, collectively reflecting obesity-associated VAT dysfunction and metabolic reprogramming as tumor progresses.

To assess the lasting impact of obesity on EV composition, we examined proteins that were consistently up- or downregulated across both early and late tumors in tumor- and VAT-EVs. In tumor-EVs, no overlapping proteins were identified, highlighting a dynamic and tumor size-specific proteomic signature. This finding reinforces the notion that tumor progression profoundly alters the tumor-EV proteome. In contrast, VAT-EVs from DIO mice exhibited greater temporal consistency: we identified 10 proteins that were commonly upregulated in both DIO-0.5 and DIO-1 groups (Table 4). These proteins are functionally associated with RNA surveillance (MOV10, UPF1), oxidative stress responses (SOD3, CRYZL2), metabolism (PRPS1, HPRT1, PLTP), and extracellular signaling (SFRP5, C1QTNF9), suggesting persistent obesity-associated alterations in adipose tissue communication that may promote tumor progression. Further, three of these proteins (CRYZL2, CYP4b1, and SOD3) were consistently elevated in late tumor-EVs, suggesting a potential obesity-associated signature that is independent of tissue source and to some extent, tumor size. We also identified 21 commonly downregulated proteins in VAT-EVs (DIO-0.5 and DIO-1), many involved in mitochondrial function, lipid metabolism, and membrane structure—reflecting sustained VAT dysfunction with obesity.

Notably, four proteins—TMEM143, MTND2, HSD11B1, and FASN—were commonly downregulated in both VAT- and tumor-EVs from DIO mice bearing early tumors. TMEM143 is a poorly characterized transmembrane protein potentially involved in mitochondrial dynamics or lipid trafficking (Nakachi et al., 2008). MTND2 is involved in electron transport chain function (Feng et al., 2012). HSD11B1 catalyzes the interconversion of active and inactive glucocorticoids, regulating local inflammatory tone and metabolic responses, and its reduced presence in EVs may reflect disrupted hormonal regulation under metabolic stress (Cooper and Stewart, 2009). Similarly, the downregulation of FASN, a central enzyme in *de novo* fatty acid synthesis (Menendez and Lupu, 2007), implies that lipogenesis is diminished in EVs during early disease stages. The consistent downregulation of these 4 proteins in VAT- and tumor-EVs from DIO mice bearing early tumors suggests a shared impairment in mitochondrial function and hormonal regulation between VAT and tumor tissues under obese conditions, which may contribute to early metabolic reprogramming of the tumor microenvironment.

Together, these findings highlight the role of obesity in shaping EV-mediated crosstalk between VAT and tumors, ultimately supporting a more aggressive and metabolically adaptable tumor microenvironment.

### Tumor- and VAT-EV Regulation of Immune Response and Metabolism

The EV proteomic profiling revealed tumor size- and obesity-associated shifts in EV composition. DIO mice had early tumor-EVs depleted in immune-related proteins and VAT-EVs enriched in mitochondrial and fatty acid oxidation proteins. In contrast, late tumor-EVs were enriched in lipid metabolism, mitochondrial adaptation, and oxidative stress resistance proteins, while VAT-EVs showed reduced mitochondrial and lipid metabolism proteins. These changes prompted functional testing of EV effects on immune responses and metabolism.

To assess how obesity-associated changes in EV composition influence immune activity and metabolism during tumor progression, we co-cultured CD8⁺ T cells with tumor- and VAT-EVs from lean and DIO mice bearing early or late tumors. First, using confocal microscopy, we confirmed cellular uptake of PKH26-labeled tumor- and VAT-EVs 24 h after co-culture (Figure 7A). Internalized EVs were observed within the cytoplasm, confined by the cell membrane boundaries.

**Figure 7:**
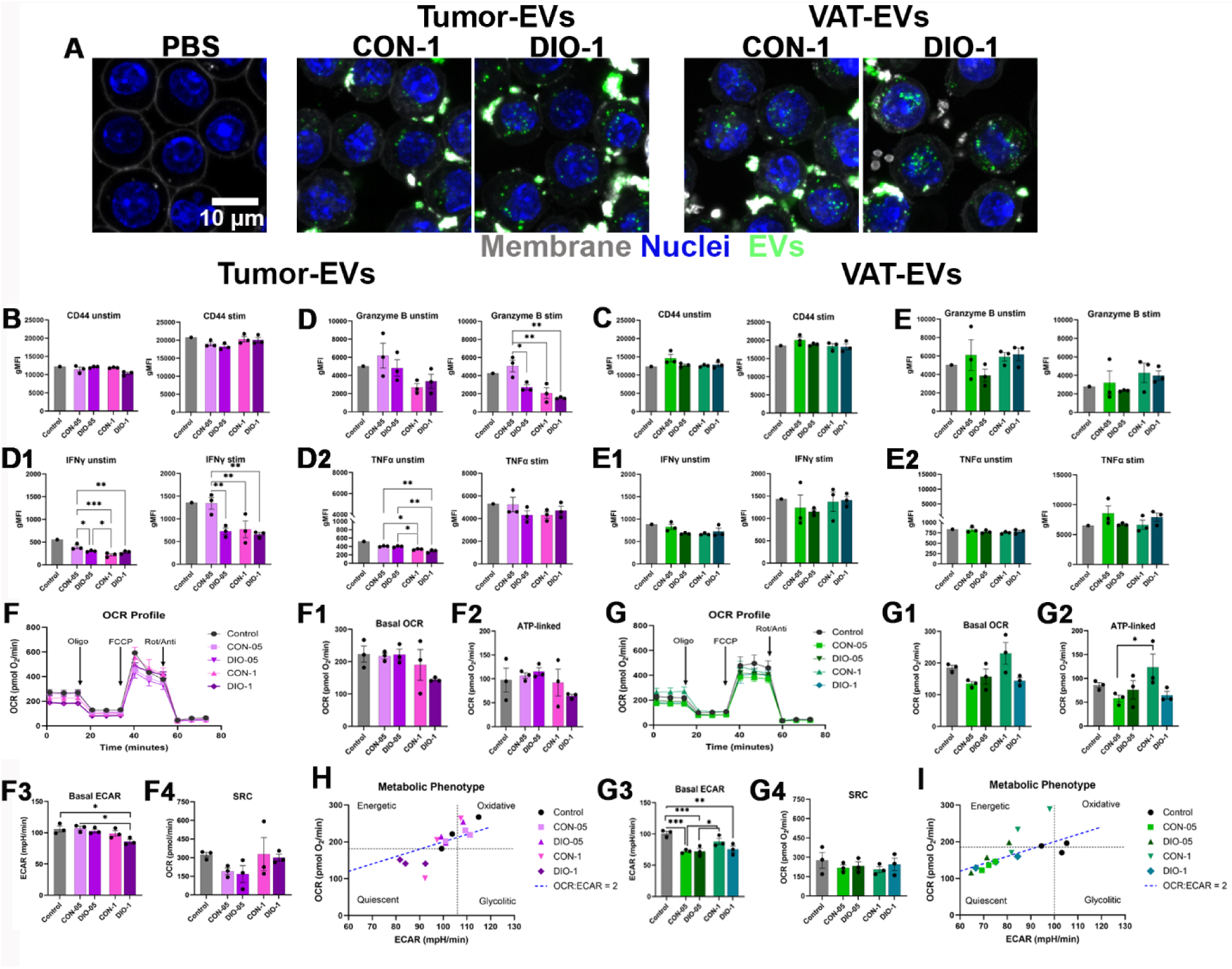
CD8⁺ T cells internalize tumor- and VAT-derived EVs, leading to altered immune activation and metabolic phenotype. **(A)** Immunofluorescence images showing uptake of tumor- and VAT-derived EVs by CD8⁺ T cells. **(B - D)** Flow cytometry analysis of CD44, granzyme B, IFNγ, and TNFα expression in unstimulated (unstim) and stimulated (stim) CD8^+^ T cells treated with 10 µg of tumor- or VAT- derived EVs. Results expressed as the geometric mean fluorescence intensity (gMFI). **(F - G)** OCR profiles and quantification of metabolic parameters, including basal oxygen consumption rate (OCR), ATP-linked respiration, basal extracellular acidification rate (ECAR) and spare respiratory capacity (SRC) following treatment with tumor- or VAT-EVs. **(H - I)** Metabolic phenotype of EV-treated CD8^+^ T cells, each point represents a biological replicate. The horizontal and vertical dashed lines represent the mean basal OCR and basal ECAR of the control samples, dividing the graph into four metabolic phenotypes: Oxidative (high OCR, low ECAR), Glycolytic (low OCR, high ECAR), Energetic (high OCR and ECAR), and Quiescent (low OCR and ECAR). The blue dashed line represents the theoretical ratio of OCR:ECAR = 2, used as a reference for metabolic balance. Data are presented as mean ± SEM from *n* = 3 biological replicates. Statistical comparisons were performed using two-way ANOVA followed by Tukey’s multiple comparisons test. Significance is indicated as: p *<* 0.05 (*\**), p < 0.01 (**), p < 0.001 (*\*\*\**), p *<* 0.0001 (****).

Next, we investigated how EVs influence CD8^+^ T cells activation and effector function. CD8⁺ T cells were treated with 50 μg of tumor- or VAT-EVs from CON and DIO mice bearing early- and late-stage tumors. CD44 expression, a marker of activation, was not significantly altered in unstimulated or anti-CD3/CD28-stimulated CD8⁺ T cells (Figure 7B and 7C), suggesting minimal activation potential of these EVs in the absence or presence of T cell receptor engagement.

To assess how EVs influence CD8⁺ T cell effector activity, we measured the levels of the protease granzyme B, and cytokines IFN-γ and TNF in both unstimulated and anti-CD3/CD28-stimulated CD8⁺ T cells following treatment with tumor- or VAT-EVs (Figure 7D-D2 and 7E-E2). In unstimulated CD8⁺ T cells, treatment with tumor- or VAT-EVs from lean mice bearing early tumors (CON-05) did not change granzyme B levels compared to vehicle control (Figure 7D and 7E). However, tumor-EVs from early (DIO-0.5), and late tumors (CON-1 and DIO-1) reduced granzyme B levels relative to early tumor-EVs from lean mice (CON-0.5). In stimulated cells, granzyme B levels were elevated in those treated with early tumor-EVs (CON-0.5), while all other tumor-EV-treated groups, including DIO-0.5 and late tumor-EVs (CON-1 and DIO-1), showed significantly lower expression (Figure 7D). In contrast, VAT-EVs had no significant effect on granzyme B levels in stimulated CD8⁺ T cells (Figure 7E). A similar pattern was observed for IFN-γ. Early tumor-EVs from lean mice (CON-0.5) had no effect in unstimulated or stimulated cells (Figure 7D1), while tumor-EVs from all other groups significantly reduced IFN-γ expression. VAT-EVs did not impact IFN-γ levels in either condition (Figure 7E1). The effects on TNF expression exhibited a distinct pattern. In unstimulated CD8⁺ T cells, tumor-EVs reduced TNF expression across all groups, with late tumor-EVs from lean and DIO mice (CON-1 and DIO-1) inducing the most pronounced decreases (Figure 7D2). VAT-EVs had no effect in unstimulated cells (Figure 7E2). In stimulated cells, TNF levels remained unchanged across all EV-treated groups. Taken together, these findings suggest that early-tumor EVs from lean mice (CON-0.5) may support CD8⁺ T cell effector functions, whereas EVs from DIO mice and late tumors impair these functions.

To assess how EVs influence CD8⁺ T cell mitochondrial function, we co-cultured cells for 48 h with tumor- or VAT-EVs purified from lean and obese mice bearing early- or late-tumors (Figure 7F, and 7G). Metabolic flux analysis showed that EV source and body weight phenotype jointly shaped CD8⁺ T cell bioenergetics. CD8⁺ T cells treated with tumor-EVs exhibited a modest reduction in oxygen consumption rate (Filippone et al.) across all respiratory states, with the most pronounced suppression observed in those treated with late tumor-EVs from DIO mice (DIO-1 group) (Figure 7F). Although basal and ATP-linked OCR trended downward in the DIO-1 group, these changes were not statistically significant (Figure 7F1 and 7F2). In contrast, basal extracellular acidification rate (ECAR), a proxy for glycolytic activity, was significantly reduced in cells treated with late tumor-EVs from DIO mice (DIO-1, Figure 7F3). Correspondingly, the metabolic phenotype mapping (Figure 7H) showed a shift toward the quiescent quadrant in CD8⁺ T cells treated with late tumor-EVs from DIO-1 mice, consistent with reduced energetic capacity.

The treatment of CD8^+^ T cells with VAT-EVs (Figure 7G) led to distinct metabolic effects. CD8D T cells treated with VAT-EVs from lean mice bearing late tumors (CON-1) showed a significant increase in ATP-linked OCR, whereas DIO-derived VAT-EVs had no significant effect on this parameter (Figure 7G2). Notably, VAT-EVs from all groups significantly suppressed basal ECAR (Figure 7G3), with the strongest reductions observed in CON-05, DIO-0.5, and DIO-1, groups. Despite minimal changes in spare respiratory capacity (SRC, Figure 7G4), metabolic phenotype plots (Figure 7I) revealed a shift toward quiescence in cells treated with VAT-EVs.

Together, these findings suggest that tumor- and VAT-EVs from DIO mice suppress oxidative metabolism in CD8⁺ T cells and reduce their ability to increase ECAR in response to mitochondrial stress, indicating impaired metabolic flexibility. In contrast, VAT-EVs from lean mice with late tumors (CON-1) may enhance mitochondrial ATP production. Overall, with obesity, late tumor-EVs and VAT-EVs appeared to compromise CD8⁺ T cell energetic fitness, potentially limiting their antitumor function.

### EVs from Tumor and Adipose Tissue Alter Energy Metabolism in Breast Cancer Cells During Obesity-Driven Tumor Progression

To evaluate how EVs derived from tumors and VAT of obese (DIO) and lean (CON) mice influence the metabolic activity of mammary cancer cells, we cultured E-Wnt (epithelial-like) and M-Wnt (mesenchymal-like) murine mammary tumor cells in EV-depleted media (ultracentrifuged FBS; UCF) and treated them with 50 µg of EVs purified from tumor or VAT during early (CON-0.5, DIO-0.5) and late (CON-1, DIO-1) tumor development. Immunofluorescence confirmed successful uptake of labeled EVs (magenta) into both E-Wnt and M-Wnt cells, with internalized vesicles localized to the cytoplasm (Figure 8A, 8B).

**Figure 8:**
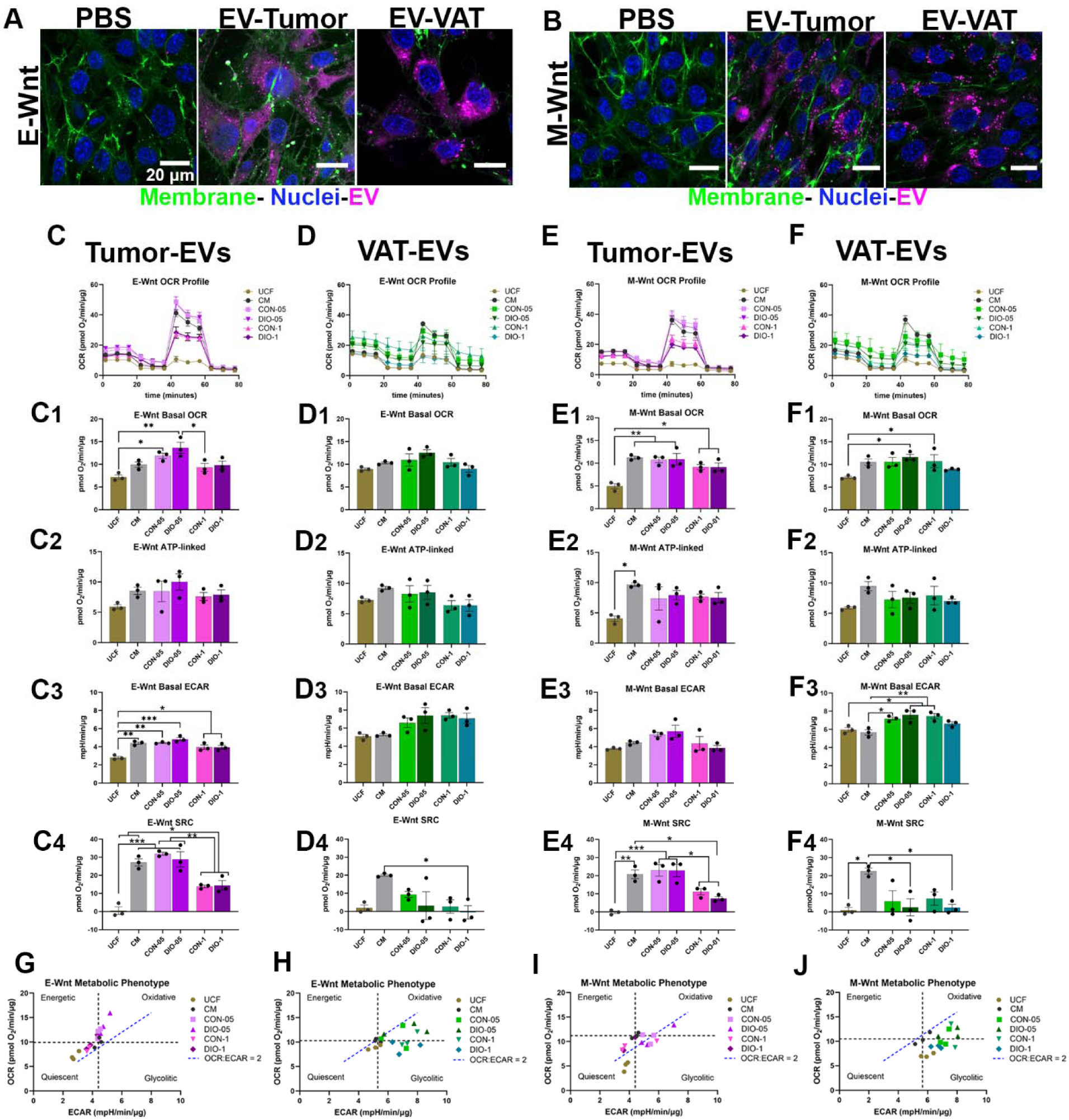
Tumor- and VAT-derived EVs from obese and lean mice differentially affect mitochondrial and glycolytic metabolism in epithelial- and mesenchymal-like breast cancer cells. E-Wnt (epithelial) and M-Wnt (mesenchymal) murine mammary tumor cells were cultured in EV-depleted media (ultracentrifuged FBS; UCF)) and treated with 50_µg of EVs isolated from tumors or VAT of control (CON) or diet-induced obese (DIO) mice during early (CON-05, DIO-05) or late (CON-1, DIO-1) stages of tumor development defined by tumor size. Cells cultured in complete media (CM) or UCF without EVs were included as control groups to assess baseline metabolic activity. **(A–B)** Immunofluorescence showing uptake of PKH26-labeled EVs (magenta) by E-Wnt (A) and M-Wnt (B) cells. **(C–F)** OCR profiles and quantification of basal OCR (C1, D1, E1, F1), ATP-linked respiration (C2, D2, E2, F2), basal ECAR (C3, D3, E3, F3), and spare respiratory capacity (SRC; C4, D4, E4, F4) in E-Wnt and M-Wnt cells treated with tumor- or VAT-derived EVs.**(G–J)** Metabolic phenotype maps showing glycolytic vs. oxidative metabolic profiles in E-Wnt (G, H) and M-Wnt (I, J) cells. The horizontal and vertical dashed lines represent the mean basal OCR and basal ECAR of the control samples, dividing the graph into four metabolic phenotypes: Oxidative (high OCR, low ECAR), Glycolytic (low OCR, high ECAR), Energetic (high OCR and ECAR), and Quiescent (low OCR and ECAR). The blue dashed line represents the theoretical ratio of OCR:ECAR = 2, used as a reference for metabolic balance. Data are presented as mean ± SEM from *n* = 3 biological replicates. Statistical comparisons were performed using two-way ANOVA followed by Tukey’s multiple comparisons test. Significance is indicated as: p *<* 0.05 (*\**), p < 0.01 (**), p < 0.001 (*\*\*\**), p *<* 0.0001 (****).

Mitochondrial stress tests revealed that tumor- and VAT-EVs induced distinct and tumor stage-dependent metabolic changes, as shown in OCR profiles (Figures 8C–F). In E-Wnt cells, tumor-EVs from early tumors (CON-0.5, DIO-0.5) increased basal OCR, whereas tumor-EVs from late tumors (CON-1, DIO-1) decreased it (Figure 8C1). VAT-EVs did not significantly alter basal OCR (Figure 8D1). Although, ATP-linked respiration, tended to be enhanced by tumor- and VAT-EVs from mice bearing early tumors and decreased by tumor and VAT-EVs from mice bearing late tumors (Figures 8C2 and 8D2), the effect was not significant.

Additionally, basal ECAR in E-Wnt cells was significantly increased by tumor-EVs from early tumors, reaching levels similar to those in complete media (CM), while EVs from late-tumors (CON-1 and DIO-1) significantly reduced ECAR (Figure 8C3). In contrast, VAT-EVs from all groups did not impact ECAR (Figure 8D3). Spare respiratory capacity (SRC) followed a similar trend in tumor-EV treated cells, with higher SRC in cells treated with EVs from early tumors (CON-0.5 and DIO-0.5) groups and reduced SRC in when treated with EVs from late tumor groups (Figure 8C4). E-Wnt cells treated with VAT-EVs from all groups, showed consistently low SRC, similar to UCF control, with a significant reduction in SRC in cells treated with VAT-EVs from DIO mice bearing late tumors compared to CM control (Figure 8D4).

In M-Wnt cells, tumor- and VAT-EVs from lean and DIO mice bearing late tumors (CON-1 and DIO-1) showed reduced basal OCR relative to the CM group (Figures 8E1 and 8F1), while ATP-linked respiration remained unchanged (Figures 8E2 and 8F2). Basal ECAR showed a nonsignificant reduction in M-Wnt cells treated with late tumor-EVs relative to the UCF and CM groups (Figure 8E3), whereas VAT-EVs from lean and DIO mice bearing early tumors (CON-0.5, DIO-05) and VAT-EVs from mice bearing late tumor (CON-1) robustly increased basal ECAR. However VAT-EVs from DIO mice bearing late tumors (DIO-1) had not significant effects on ECAR (Figure 8F3). Notably, SRC was significantly reduced in M-Wnt cells treated with tumor-EVs from late tumors (CON-1, DIO-1) and VAT-EVs from DIO mice bearing late tumor, relative to CM control (Figures 8E4 and 8F4), indicating a marked impairment in mitochondrial reserve capacity in the more aggressive mesenchymal cell population. Metabolic phenotype mapping (Figures 8G–J) further illustrated these findings. In E-Wnt cells, treatment with tumor-EVs from early tumors promoted an oxidative phenotype, while late tumor-EVs (CON-1, DIO-1) induced a shift toward a quiescence (Figure 8G). In M-Wnt cells, early tumor-EVs increased glycolytic activity, whereas late tumor-EVs suppressed metabolic activity torwards a quiescence (Figure 8I). In contrast, VAT-EVs maintained an oxidative phenotype in both E-Wnt and M-Wnt cells, but shifted the cells toward a more glycolytic profile, particularly when treated with VAT-EVs from lean and DIO mice bearing late tumors (CON-1 and DIO-1, Figures 8H and 8J). Together, these data suggest that EVs derived from tumors and VAT dynamically reprogram the metabolic activity of breast cancer cells in a manner dependent on tumor size, EV source, and tumor cell phenotype. Obesity-associated EVs from late-stage tumors exert the most pronounced suppressive effects on mitochondrial function—particularly in mesenchymal-like M-Wnt cells—while VAT-derived EVs preferentially shift cells toward a glycolytic profile, highlighting the multifaceted role of EVs in shaping the tumor metabolic landscape during obesity-driven cancer progression.

## Discussion

EVs are increasingly recognized as key mediators of intercellular communication in cancer, integrating signals related to metabolism, immune regulation, and tissue remodeling. While prior studies have cataloged EV proteomes across tumors and circulation, and stromal compartments —identifying candidate biomarkers and functional pathways linked to metastasis and disease severity (Hoshino et al., 2020, Leung et al., 2021, Zhang et al., 2024b, Mallawaaratchy et al., 2017)— how obesity and tumor progression jointly shape EV composition across distinct tissue sources, and how these changes translate into functional immune and metabolic consequences, has remained unclear, particularly in obesity-driven TNBC.

In this study, we use tumor volume as a proxy for tumor progression. Tumor size is a well-established clinical indicator of disease stage in human breast cancer, with larger tumors associated with more aggressive behavior (Amin et al., 2017). Although histological characterization was not performed here, prior work using this same model has shown that larger tumors are more undifferentiated and metastatic (O’Flanagan et al., 2017). This size-based stratification in a well stablished preclinical model of DIO and TNBC enabled us to begin disentangling the independent and interacting effects of tumor progression and obesity on EV proteomic remodeling.

Using this framework, we show that tumor progression and DIO impose distinct yet interacting effects on the proteome of EVs. Tumor-EVs and VAT-EVs exhibited divergent proteomic trajectories across tumor stages, supporting the concept that EVs composition reflects information about both tissue origin and microenvironmental cues (Hoshino et al., 2015, Brzozowski et al., 2018, Wortzel et al., 2019). Tumor progression primarily shaped the tumor-EV proteome, whereas obesity exerted a dominant influence on VAT-EVs, consistent with prior reports of tissue-specific EV adaptation during metabolic disease and cancer (Le Lay and Scherer, 2025, Zhang et al., 2026, Xavier et al., 2020, Monoe et al., 2025)

Across tumor stages within the same dietary group, tumor-EVs evolved from profiles enriched in cytoskeletal organization and nucleotide metabolism in early tumors toward signatures dominated by metabolic rewiring, stress adaptation, and ECM remodeling in late tumors. Obesity amplified these late tumor features, with tumor-EVs from DIO mice enriched in proteins involved in lipid metabolism, oxidative stress resistance, immune modulation, and ECM remodeling—hallmarks associated with aggressive tumor phenotypes(Hanahan, 2022, Li et al., 2025). These findings align with prior reports showing that EV composition varies with tissue origin, tumor aggressiveness, and microenvironmental context (Hoshino et al., 2015, Brzozowski et al., 2018, Wortzel et al., 2019)

VAT-EVs displayed a distinct and temporally divergent trajectory. In lean mice bearing early tumors, VAT-EVs were enriched in immune and metabolic regulators, suggesting a metabolically active adipose environment capable of supporting local and systemic homeostasis. In contrast, VAT-EVs from obese mice at early stages showed enrichment in ECM remodeling and signaling proteins, consistent with adipose tissue inflammation and stromal remodeling in obesity. As tumors progressed, VAT-EVs from DIO mice shifted toward stress-response markers accompanied by loss of mitochondrial and lipid metabolism proteins, reflecting progressive adipose tissue dysfunction—a hallmark of chronic obesity (Le Lay et al., 2021, Crewe et al., 2021, Malaguarnera et al., 2025). Together, these findings indicate that obesity not only alters tumor- EV signaling but also profoundly reprograms VAT-EVs in a manner that evolves with tumor burden.

Comparison of EV composition between DIO and control mice within the same tumor stage further revealed obesity-specific EV signatures with potential functional consequences. Early-stage tumor-EVs from DIO mice were enriched in immune signaling molecules, RNA processing proteins, and regulators of microenvironmental remodeling, suggesting that obesity-associated tumor-EVs may contribute to priming the tumor niche and modulating host responses during early tumor establishment (Zhong et al., 2025). In contrast, tumor-EVs from DIO mice bearing late tumors emphasized lipid metabolism, oxidative stress regulation, and ECM remodeling, including enzymes involved in fatty acid catabolism (e.g., MGLL, LIPE, ACSL1), mitochondrial redox balance (e.g., SOD3, ALDH6A1), and structural remodeling (e.g., COL1A1, COL5A2). The presence of metabolic enzymes such as pyruvate carboxylase (PC), which sustains tricarboxylic acid cycle activity under hypoxic conditions, further implicates EVs as vehicles for metabolic adaptation supporting tumor survival and progression (Shinde et al., 2018, Kiesel et al., 2021)

Beyond diet- and stage-specific changes, we identified overlapping EV proteins that may represent conserved responses to tumor progression independent of obesity. Recurrent upregulation of proteins involved in ECM organization, (TIH4, ITGA2B) (Zhang et al., 2024a, Podstawski et al., 2022), stress response (HSPB1, HSPA1A) (Kishi et al., 2016, Azzazene et al., 2013), and metabolic and immune signaling regulators (PROCR, GPLD1, ADPRH, and ENPP1) (Cao et al., 2023, Kishi et al., 2016, Zhang et al., 2021, Huang et al., 2024), suggests the emergence of a core EV signature supporting processes associated with tumor aggressiveness (Hanahan, 2022). Similarly, the consistent enrichment of EIF2AK3 (PERK) in VAT-EVs across dietary conditions highlights a potential mechanism by which adipose tissue communicates stress-adaptive signals to the tumor microenvironment. PERK-mediated activation of the eIF2α/ATF4 axis has been linked to therapy resistance and metastasis in TNBC (Feng et al., 2017, Bai et al., 2021, Calvo et al., 2023), and adipose- EVs have been shown to induce endoplasmic reticulum stress in recipient cells (Le Lay et al., 2021).

Persistent obesity-associated EV signatures shared between tumor- and VAT-EVs—including CRYZL2, SOD3, and CYP4B1—suggests coordinated EV-mediated communication between adipose tissue and tumors under obese conditions. These proteins participate in redox regulation, lipid metabolism, and xenobiotic processing, collectively shaping a pro-tumorigenic and immunosuppressive microenvironment (Fukai and Ushio-Fukai, 2011, Carmona-Rodríguez et al., 2020, Sun et al., 2023).

Functionally, obesity-driven EV remodeling was linked to impaired antitumor immunity. EVs from obese mice — particularly tumor-EVs from late tumors — suppressed CD8D T cell mitochondrial activity and effector cytokine production, highlighting EV-mediated metabolic reprogramming as a mechanism contributing to immune dysfunction in obesity. Tumor-EVs are known to inhibit T cell function through delivery of immunosuppressive and metabolically disruptive cargo (Whiteside, 2016, Poggio et al., 2019) counteracting the metabolic flexibility required for robust effector function (Scharping et al., 2016, O’Sullivan et al., 2014). In contrast, VAT-EVs from lean mice bearing tumor enhanced mitochondrial ATP production, suggesting that VAT-EVs can support immunometabolic fitness under non-obese conditions — a capacity that is lost with obesity. These findings extend existing models of obesity-driven immune suppression by implicating EVs as active regulators of T cell metabolic flexibility (Ringel et al., 2020, Dyck et al., 2022).

EV-mediated metabolic reprogramming also extended to tumor cells in a stage- and phenotype-dependent manner. Tumor-EVs from early tumors promoted energetically active phenotypes in epithelial-like tumor cells (E-Wnt) consistent with reports that tumor EVs enhance mitochondrial respiration and metabolic fitness to support growth (Gelsomino et al., 2022). In contrast, EVs from late tumors suppressed both oxidative and glycolytic metabolism, particularly in mesenchymal-like cells (M-Wnt) suggesting a shift toward metabolic restraint or stress adaptation associated with aggressive tumor state (Hanahan, 2022). VAT-EVs exerted complementary effects, broadly enhancing glycolytic activity while, under obese conditions and late tumor, further constraining mitochondrial reserve capacity.

Together, these findings support a unified model in which obesity amplifies EV-mediated communication between tumors, adipose tissue, and immune cells, promoting metabolic reprogramming and immune suppression during TNBC progression. While this study focused on tumor- and VAT-derived EVs, future work incorporating circulating EVs, longitudinal sampling, and spatial tumor profiling will be essential to fully resolve EV-mediated inter-organ signaling in obesity-driven cancer.

Targeting EV-dependent metabolic and immune pathways may represent a promising therapeutic strategy in obesity-associated TNBC.

## Acknowledgements

We thanks the UNC metabolomics and Proteomics (MAP) core.

## Supplemental data

This article contains supplemental data

## Author contributions

X.B.M. conceptualization; X.B.M., E.J.G., L.A.R., E.R and Y.A. formal analysis; X.B.M, E.J.G., L.A.R. Y.A. investigation; S.D.H., D.T. and N. McG. resources; X.B.M and E.J.G. writing – original draft; X.B.M, and S.D.H writing – review & editing; X.B.M. visualization; X.B.M. supervision; X.B.M. and E. R project administration; X.B.M., S.D.H, and D.T. funding acquisition. All authors reviewed the final manuscript.

