## Supplementary Figures for "Obesity and Tumor Development Reprogram the Proteome and Metabolic Effects of Adipose- and Tumor-Derived Extracellular Vesicles"

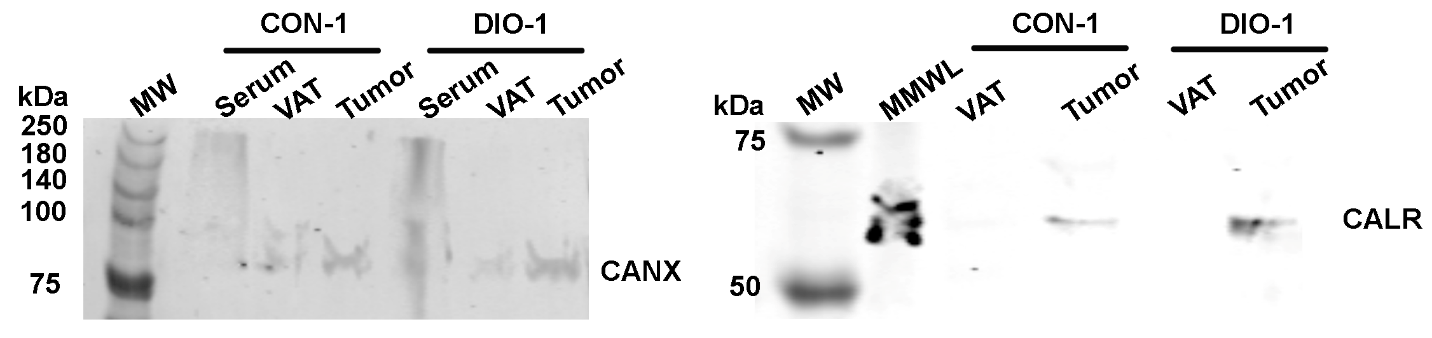

**Supplementary Figure 1: Western blot analysis of EVs.**

Representative western blot showing the presence of calnexin (CANX) and calreticulin (CALR) in EV preparations from serum, VAT and tumors purified from CON or DIO mice. Calnexin and calreticulin, both endoplasmic reticulum (ER) resident proteins, were detected in cell lysates of metM-Wnt^lung^ cells (MMWL) but were markedly reduced in EV samples, consistent with their use as negative markers for EV purity. Equal amounts of protein were loaded for each condition.

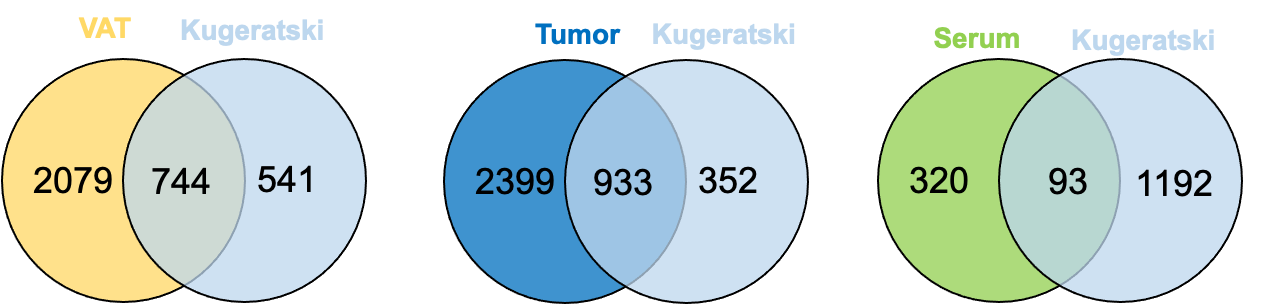

**Supplementary Figure 2:. Overlap between study-derived EV proteins and the Kugeratski EV proteome.**Venn diagrams illustrating the overlap between proteins identified in this study and the reference list of 1,285 EV-associated genes reported by Kugeratski et al. (2021), which represent 1,212 proteins commonly found across various EV subtypes and isolation methods. Of the global proteome identified in VAT-derived EVs, 744 (58%) genes overlapped with the Kugeratski dataset. Tumor-derived EVs showed the highest overlap, with 933 (72%) matching genes. In contrast, only 93 (7%) of the proteins identified in serum-derived EVs were shared with the reference list. These results highlight tissue-specific differences in EV protein composition and alignment with established EV markers.
